# Mapping the aggregate g-ratio of white matter tracts using multi-modal MRI

**DOI:** 10.1101/2024.09.25.614579

**Authors:** Wen Da Lu, Mark C. Nelson, Ilana R. Leppert, Jennifer S.W. Campbell, Simona Schiavi, G. Bruce Pike, Christopher D. Rowley, Alessandro Daducci, Christine L. Tardif

## Abstract

The g-ratio of a myelinated axon is defined as the ratio of the inner-to-outer diameter of the myelin sheath and modulates conduction speed of action potentials along axons. This g-ratio can be mapped *in vivo* at the macroscopic scale across the entire human brain using multi-modal MRI and sampled along white matter streamlines reconstructed from diffusion-weighted images to derive the g-ratio of a white matter tract. This tractometry approach has shown spatiotemporal variations in myelin g-ratio across white matter tracts and networks. However, tractometry is biased by partial volume effects where voxels contain multiple fiber populations. To address this limitation, we used the Convex Optimization Modeling for Microstructure-Informed Tractography (COMMIT) framework to derive tract-specific axonal and myelin volumes, which are used to compute the tract-specific aggregate g-ratio. We compare our novel COMMIT-based tract-specific g-ratio mapping approach to conventional tractometry in a group of 10 healthy adults. Our findings demonstrate that the tract-specific g-ratio mapping approach preserves the overall spatial distribution observed in tractometry and enhances contrast between tracts. Additionally, our scan-rescan data shows high repeatability for medium to large caliber tracts. We show that short and large caliber tracts have a lower g-ratio, whereas tractometry results show the opposite trends. This technique advances tract-specific analysis by reducing biases introduced by the complex network of crossing white matter fibers.

## 1 Introduction

The g-ratio is a measure of the thickness of the myelin sheath relative to the caliber of the axon and is calculated as the ratio between the inner and outer diameters of the myelin sheath. Myelin plays a vital role in promoting rapid and efficient communication between neurons in the brain through mechanisms such as saltatory conduction, metabolic support and shorter refractory periods (Khelfaoui et al., 2024). Myelin plasticity is also a mechanism to modulate the precise timing of signals within brain networks to support learning and higher cognitive functions (Xin & Chan, 2020). The myelin g-ratio is one of the primary modulators of axonal conduction velocity, although other features including inter-node length, myelin periodicity, and axon caliber also contribute (Drakesmith et al., 2019). Myelination is a protracted process that peaks around mid-life (Slater et al., 2019), although myelin remains adaptive throughout the lifespan (Xin & Chan, 2020). Demyelination and atypical myelination have been observed in several neurological and psychiatric disorders such as multiple sclerosis (Frohman et al., 2006) and schizophrenia (Davis et al., 2003). Measuring the g-ratio of brain tissue *in vivo* is thus critical to improve our understanding of how myelination supports brain function during neurodevelopment, aging, and learning, and how dysmyelination alters brain function and behaviour.

Histological studies have provided foundational insights into the g-ratio. During human neurodevelopment, axon growth initially surpasses myelination, resulting in a declining g-ratio as myelination progresses (Schröder et al., 1988). The g-ratio of myelinated axons varies with axon caliber, being approximately 0.6 for axons smaller than 5 µm in diameter and greater than 0.6 for axons larger than 5 µm in diameter (Graf von Keyserlingk & Schramm, 1984). These findings align with the concept of an optimal g-ratio, which balances the spatial constraints of the central nervous system and cellular energetics, generally falling within the range of 0.6–0.8 (Chomiak & Hu, 2009). These histological studies were labor-intensive and limited by small fields of view and specific cutting planes (perpendicular to the axon). *In vivo* measurement techniques are needed to overcome these limitations and further elucidate the role of g-ratio in brain function and pathology.

We can non-invasively estimate the g-ratio *in vivo* across the whole brain using multi-modal, quantitative magnetic resonance imaging (MRI). Stikov, Campbell et al. showed that the aggregate g-ratio of each voxel can be computed from MRI markers of myelin volume fraction (MVF) and intra-axonal volume fraction (AVF) using a simple geometric model, without the need to acquire new MR contrasts (Stikov et al., 2015). The aggregate g-ratio of a voxel represents a weighted average of the g-ratio of each axon where larger axons will have a greater weight. In this article, we will refer to the “aggregate g-ratio” simply as the “g-ratio”. MRI-based g-ratio mapping methods and applications are reviewed in (Campbell et al., 2018) and (Mohammadi & Callaghan, 2021).

The g-ratio of specific white matter tracts can be estimated using a tractometry pipeline (Bells et al., 2011) where the g-ratio map is projected onto streamlines reconstructed from diffusion-weighted MRI using tractography. The mean or median g-ratio value can then be computed across a bundle of streamlines to create a g-ratio tract profile (Yeatman et al., 2012). The mean or median g-ratio can also be computed along the length of the bundle of streamlines representing a white matter tract (Slater et al., 2019). The tracts can be defined using regions of interest for inclusion or exclusion of streamlines or using tract atlases (Radwan et al., 2022). Alternatively, a whole brain structural connectome can be reconstructed where each edge or tract corresponds to the bundle of streamlines connecting two nodes of a cortical and deep grey matter parcellation (Kamagata et al., 2019). However, the true anatomical specificity of tractometry is limited due to the complex wiring geometry of the brain. It is estimated that 60-90% of white matter voxels contain multiple fiber populations at typical diffusion imaging resolutions (Jeurissen et al., 2013), where each fiber may have a different g-ratio. This partial volume effect introduces a bias that is compounded as the g-ratio is averaged along a white matter tract, reducing the specificity of the tract g-ratio measurement and potentially obscuring subtle differences between tracts or individuals.

Despite these limitations, tractometry remains a powerful tool for studying the g-ratio of white matter tracts and networks. Slater and colleagues (Slater et al., 2019) show that g-ratio measurements of white matter tracts in 801 participants aged 7-84 years conformed most closely to a quadratic aging model, with the lowest g-ratios around the third decade of life followed by an increase, indicating thinner myelin sheaths with advancing age. Multiple groups (Berman et al., 2019; Clark et al., 2022; Mancini et al., 2021) also leveraged tract g-ratio and tract axon diameter data to estimate conduction velocity and delays. Additionally, tractometry has been used to detect significant increases in brain networks g-ratios of multiple sclerosis patients, particularly in tracts within the motor, the somatosensory, the visual, and the limbic regions (Kamagata et al., 2019).

To minimize partial volume effects and improve the anatomical specificity tract g-ratio measurements, we use the Convex Optimization Modeling for Microstructure Informed Tractography (COMMIT) framework (Daducci et al., 2014). The original COMMIT implementation estimates the effective intra-axonal cross-sectional area of the biological fibers represented by each streamline in a tractogram, using multi-shell diffusion data. It assumes that the microstructural property, in this case the intra-axonal cross-sectional area, is constant along each streamline and uses whole-brain data for signal fitting. The COMMIT framework can be adapted to other MRI modalities, provided that the signal contributions from the different tracts to a voxel are additive. For instance, COMMIT was used to estimate the myelin volume of white matter tracts from a tractogram and a myelin volume fraction map acquired separately (Schiavi et al., 2022). The COMMIT framework has also been applied to multi-contrast encoding protocols to estimate other tract-specific MR parameters: tract-specific intra-axonal T2 times (Barakovic et al., 2021) and tract-specific magnetization transfer ratio (Leppert et al., 2023).

In this study, we use the COMMIT framework to calculate tract-specific g-ratio. By combining the tract intra-axonal volumes and myelin volumes estimated using COMMIT, tract-specific aggregate g-ratio can be computed. We map the tract-specific g-ratio in 10 healthy participants and compare its spatial distribution, scan-rescan repeatability, and inter-subject variability to state-of-the-art g-ratio tractometry.

## 2 Methods

### MRI data acquisition

This study was approved by the Research Ethics Board of McGill University Health Centre, Canada, and all participants provided written informed consent. Ten participants (6 males, 29.2± 6.29 years old) were recruited, all of whom reported no prior history of neurological or psychiatric disorders. Imaging was performed on a Siemens Prisma-Fit 3 Tesla scanner using a 32-channel head coil at the McConnell Brain Imaging Centre of the Montreal Neurological Institute. All 10 participants were rescanned within a 3-week interval.

The MRI acquisition parameters are detailed in Table 1. A 1-mm isotropic T1-weighted anatomical image was acquired using MPRAGE for registration to an atlas, brain tissue segmentation, deep gray matter structure segmentation, and cortical surface extraction. 2.6-mm isotropic diffusion weighted imaging (DWI) data were acquired using a multi-shell 2D diffusion-weighted spin echo EPI sequence for whole brain tractography and intra-cellular volume fraction mapping. Magnetization transfer saturation (MTsat) maps were computed from an MT-weighted gradient echo (GRE) image, the T1 relaxation time and S0 equilibrium signal maps derived from the MP2RAGE sequence (Marques et al., 2010), and a B1^+^ map using a preconditioning pulse (Chung et al., 2010), as previously described (Rowley et al., 2024). All sequences used the Siemens *Prescan Normalize* option to generate maps with minimal receive B1^-^ field bias. A gain factor of 2.5 was applied to the S0 maps from the MP2RAGE protocols to match the receiver gain of the MT-weighted GRE image prior to computing MTsat.

**Table 1:** MRI acquisition parameters. TR = repetition time, TE = echo time, TI = inversion time, TF = turbofactor, HSn = hyperbolic secant (n=1).

| Parameter | MPRAGE | 2D diffusion-weighted spin echo EPI | MT-weighted GRE | MP2RAGE | B1<sup>+</sup> map |
| --- | --- | --- | --- | --- | --- |
| TR (ms) | 2300 | 3000 | 27 | 5000 | 20 000 |
| TE (ms) | 2.98 | 57 | 2.76 | 2.66 | 2.22 |
| Flip Angle (degrees) | 8 | 90 | 6 | 4/5 | 8 |
| TI (ms) | 900 |  | 940/2830 |  |  |
| Acceleration | GRAPPA 2 | multiband acceleration factor = 3, | 6/8 Partial Fourier GRAPPA 2. | 4.6x Upsampling, density = 0.5, | TF = 96 |
| | | 6/8 Partial Fourier | TF = 1 | jitter radius = 1.2,<br>20 iterations,<br>$6e^{-4}$ reg. TF = 175 | |
| Voxel Size<br>(mm) | 1×1×1 | 2.6×2.6×2.6 | 1×1×1 | 1×1×1 | 2.5×2.5×3 |
| Diffusion-<br>parameters | | 104 diffusion directions:<br>10 at $b = 300 \text{ s/mm}^2$,<br>30 at $b = 1,000 \text{ s/mm}^2$,<br>64 at $b = 2,000 \text{ s/mm}^2$,<br>6 $b = 0 \text{ s/mm}^2$ images. | | | |
| Pulse-<br>parameters | | | 12 ms Gaussian,<br>$\Delta = 2 \text{ kHz}$<br>$B1_{rms} = 3.26 \text{ } \mu\text{T}$ | HSn Inversion<br>pulse: 10.24 ms,<br>$B1_{rms} = 9.42 \text{ } \mu\text{T}$ | |

### MR data preprocessing

The multi-modal MRI pre-processing pipeline *micapipe* (v0.1.5) (Cruces et al., 2022) was adapted to preprocess the anatomical and the diffusion data and is described in more detail below.

#### Structural MRI analysis

The T1-weighted MPRAGE image was corrected for intensity nonuniformity, intensity normalized, skull stripped, and cortically segmented using Freesurfer (v6.0) (Dale et al., 1999; Fischl, Sereno, & Dale, 1999; Fischl, Sereno, Tootell, et al., 1999). The subcortical segmentations were performed with FSL (v6.0.3) FIRST (Patenaude et al., 2011) and the tissue types were classified using FSL FAST (Zhang et al., 2001). A five-tissue-type image segmentation was generated for anatomically constrained tractography (Smith et al., 2012).

#### DWI preprocessing and tractography

Diffusion preprocessing was performed in native DWI space using MRtrix3 (v3.0.3) (Tournier et al., 2019) and proceeded in the following sequence: (1) image denoising (Cordero-Grande et al., 2019; Veraart, Fieremans, et al., 2016; Veraart, Novikov, et al., 2016), (2) Gibbs ringing correction (Kellner et al., 2016), (3) four b = 0 s/mm^2^ volumes with reverse phase encoding were used to correct for susceptibility distortion, head motion, and eddy currents via FSL’s eddy and TOPUP tools (Andersson et al., 2003; Andersson & Sotiropoulos, 2016; Skare & Bammer, 2010; Smith et al., 2004) and (4) B1 bias-field correction (Tustison et al., 2010). The preprocessed DWI was upsampled to 1-mm to match the resolution of the T1-weighted image. The upsampled preprocessed data were used to estimate multi-shell and multi-tissue response functions for constrained spherical deconvolution (Jeurissen et al., 2014; Tournier et al., 2004) followed by intensity normalization (Dhollander et al., 2021; Raffelt et al., 2017). The T1-weighted anatomical images were nonlinearly registered to DWI space using ANTs (Avants et al., 2008).

Anatomically constrained tractography was performed on the normalized white matter fiber orientation distributions (FOD) image using the probabilistic algorithm iFOD2 (Smith et al., 2012; Tournier et al., 2010). A tractogram of 3 million streamlines was generated using the following parameters: min/max length = 10/400, max angle = 22.5, step = 0.5, cutoff = 0.06, backtrack, crop_at_gmwmi (gray-matter-white-matter interface).

#### Volumetric AVF, MVF, and g-ratio maps

The MP2RAGE-based S0 and T1 maps were fit using a dictionary matching approach that incorporates the B1^+^ map (https://github.com/JosePMarques/MP2RAGE-related-scripts) to extract ΔB1^+^ corrected S0 and T1 maps (Marques & Gruetter, 2013). The MTsat maps were computed using (Helms et al., 2008):

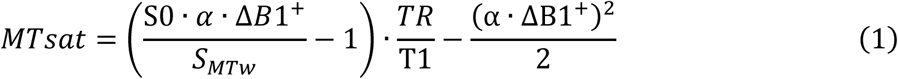

where α is the excitation flip angle in the MT-weighted image, S_MTw_ is the MT-weighted signal, and TR is the repetition time of the MT-weighted sequence. A model-based correction was used to correct for ΔB1^+^ (Rowley et al., 2021).

The diffusion data was analyzed using the NODDI model (Zhang et al., 2012) in the AMICO framework (Daducci et al., 2015) to map the intra-cellular volume fraction (ICVF) and isotropic volume fraction (ISOVF). Assuming a group average g-ratio of 0.7 in the splenium, the MVF of the splenium was estimated using equations 2-3 (Mohammadi et al., 2015; Stikov et al., 2015)

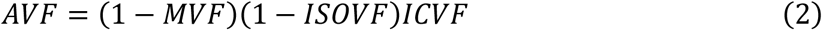

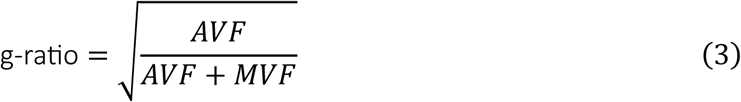

and is then employed to calculate the calibration factor α_*calib*_ below to derive the MVF map across the entire white matter.

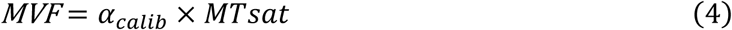

After calibration, the g-ratio is computed across the whole brain using the MVF and ICVF maps and equations 2 and 3.

### Tract-specific g-ratio estimates

The pipelines for tractometry and tract-specific g-ratio mapping are summarized in Figure 1. The tractogram generated from MRTrix3 undergoes a series of processing steps to derive tract-specific g-ratio estimates. First, the tractogram is filtered using COMMIT (v2.1, StickZeppelinBall model, parallel diffusivity of the stick and zeppelin *D*_∥_ = 1.7E-3 mm^2^/s, perpendicular diffusivity of the zeppelin *D*_⊥_ = 0.51E-3 mm^2^/s, isotropic diffusivity to account for partial voluming with gray matter and cerebral spinal fluids *D*_*iso*_= 1.7E-3, 3.0E-3 mm^2^/s) to remove implausible streamlines and to generate a volumetric map of tract ICVF.

**Figure 1:**
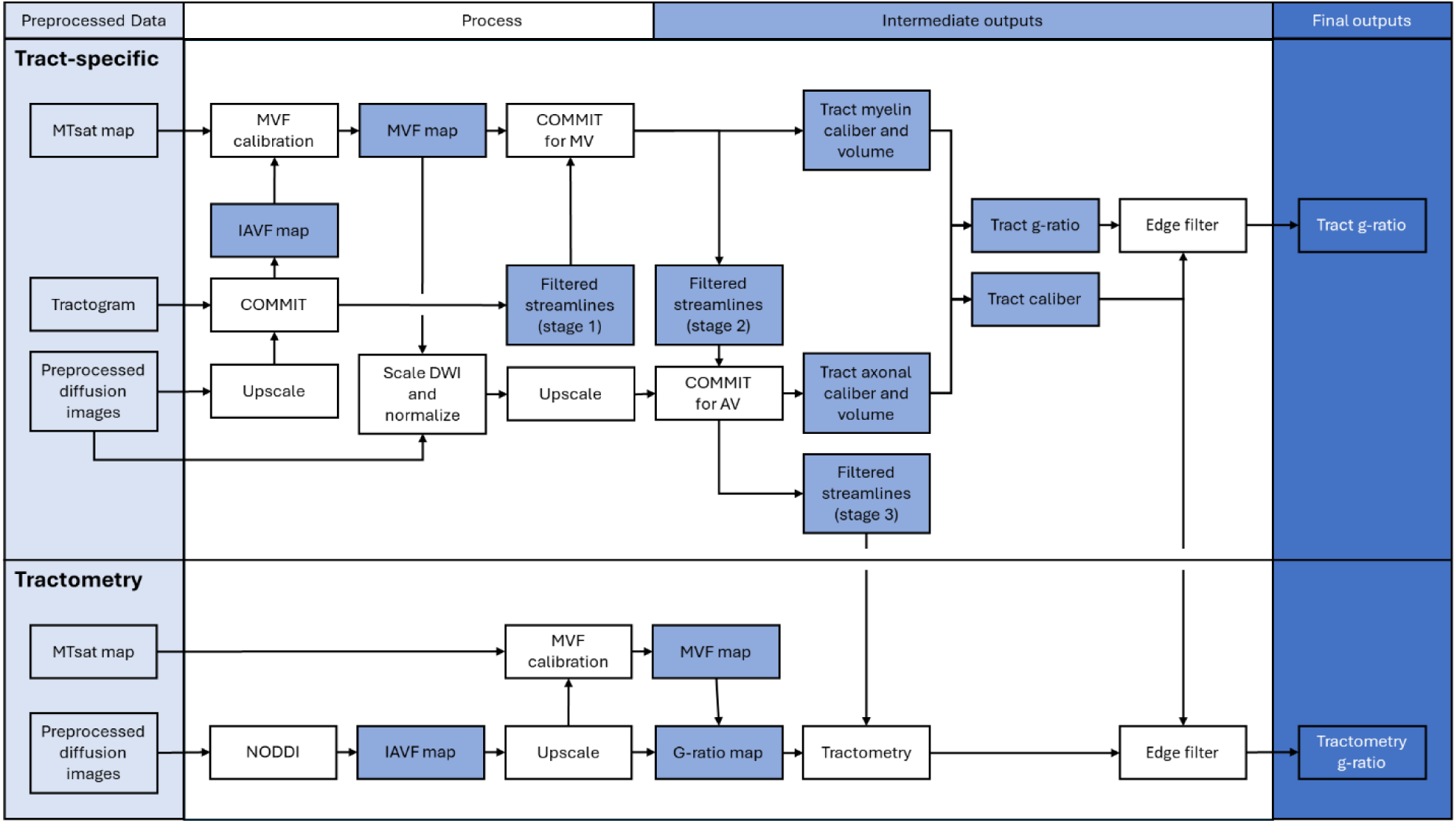
Flowchart of the processing pipeline for calculating tract-specific (above) and tractometry (below) g-ratio from multi-modal, quantitative MRI.

Assuming a splenium g-ratio of 0.7, the ICVF map generated by COMMIT is used to calibrate the MTsat map to obtain a MVF map. The MVF map and the filtered streamlines were processed through COMMIT to obtain the myelin cross-sectional area for each streamline, which was then multiplied by the streamline length to calculate the myelin volume (MV). The streamlines with a weight of 0 were removed, as they correspond to tracts deemed implausible by COMMIT.

The diffusion signal was adjusted to account for the myelin signal contribution at the voxel level. Specifically, the diffusion signal *S*(*q*) was scaled by a factor of (1−MVF), and then normalized by *S*(*q* = 0) (i.e., the b0 images) at the native DWI resolution. Consequently, the “doNormalizeSignal” option within COMMIT was set to “False”. The scaled diffusion images were upsampled to match the resolution of the T1-weighted image and input into COMMIT to compute each streamline’s axonal cross-sectional area, calculated as the sum each streamline’s *F*_*i*_ restricted diffusion (or intra-cellular) compartment signal contribution 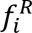 across all the voxels it traverses using the equation below

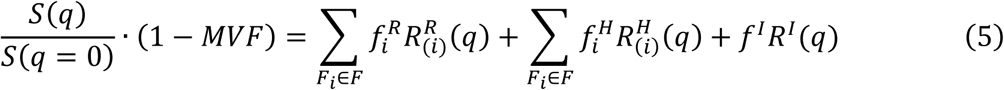

where 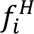 accounts for the hindered diffusion compartment signal contribution in the direction of *F*_*i*_. Additionally, the 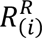 refers to the rotated version of the streamline’s response function, scaled by its length within the voxel. A similar scaling applies to the 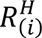 response function. Finally, the isotropic contribution is described by the signal profile *R*^*I*^(*q*), with *f*^*I*^representing the corresponding volume fraction. The cross-sectional area is then multiplied by the streamline’s length to obtain the axonal volume (AV).

MV- and AV-annotated connectivity matrices are generated from the filtered streamlines by summing their MV and AV weights. This process is performed using two cortical parcellation resolutions, Schaefer-200 and Schaefer-400 (Schaefer et al., 2017), combined with seven bilateral subcortical regions, including the amygdala, thalamus, caudate, nucleus accumbens, putamen, hippocampus, and pallidum (totaling 214 and 414 nodes, respectively). Lastly, the tract-specific g-ratio connectivity matrix is calculated from the MV- and AV-connectivity matrices using equation 3.

### Tractometry g-ratio estimates

For tractometry, we used the same filtered tractogram as for the tract-specific pipeline, allowing for an equitable comparison between the two techniques. The g-ratio of each streamline was determined by calculating the median g-ratio value sampled along the streamline’s length from the volumetric g-ratio map, as the median is more robust against outliers (Boshkovski et al., 2021). Subsequently, each edge in the connectivity matrix was computed as the mean g-ratio across all the streamlines connecting the respective node pair.

### Connectivity matrix filtering

We filtered the edges of the tract-specific connectivity matrix to improve data quality and consistency. First, we removed edges corresponding to the bottom 80% of tract calibers, calculated as the sum of their axonal and myelin cross-sectional areas, across the dataset. This step was performed to make the density of the tract-specific connectivity matrix comparable to the values reported by Luppi & Stamatakis (Luppi & Stamatakis, 2021), who used the HCP-1021 template (Van Essen et al., 2013). This process also removed small-caliber tracts with low scan-rescan repeatability. Second, we applied a 50% consensus filter across individuals, retaining only those edges present in at least half of the participants.

We conducted scan-rescan repeatability assessments of the tract-specific g-ratio technique and performed a comparative analysis with tractometry. We also explored the relationship between tract length, caliber, and g-ratio for both techniques.

## 3 Results

### Tract filtering is critical for accurate and repeatable tract-specific g-ratio estimates

The scan-rescan repeatability results of the connectivity matrix g-ratio estimates derived from the Schaefer-200 parcellation are displayed in Figure 2. While there is a concentration of points along the line of perfect fit, indicating some agreement, the unfiltered results demonstrate overall poor repeatability as evidenced by the low intra-class correlation (ICC) of 0.197 and wide limits of agreement in the Bland-Altman plot. The tracts with low repeatability correspond to those of smaller caliber (shown in Figure S1). We removed the bottom 80% of tracts based on their caliber which led to an increase in ICC to 0.720, demonstrating that g-ratio estimates of larger caliber tracts are more repeatable. Additionally, this process reduces the connectivity matrix density from 49.40% to 10.17%. Implementing a 50% consensus filter across participants further increases repeatability (ICC = 0.735) and decreases matrix density decreasing to 8.40%. This final density is similar to the 7.40% density of the HCP-1021 template. Similar trends were observed when examining the Schaefer-400 parcellation, as shown in Figure S2, with increased reproducibility following the percentile and consensus filters.

**Figure 2:**
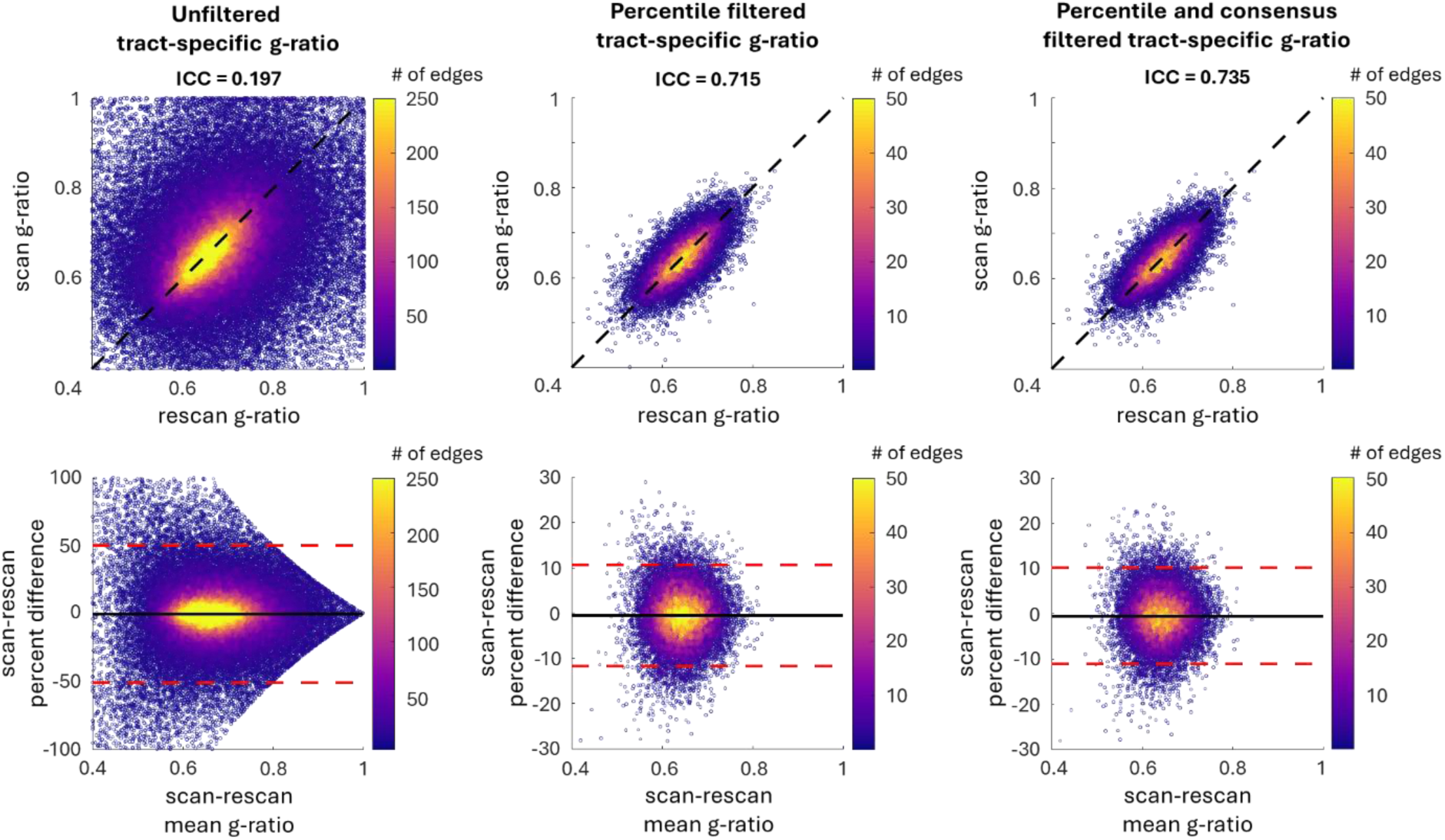
Scan-rescan repeatability of tract-specific g-ratio for the Schaefer 200 connectivity matrix. Correlation (top row) and Bland-Altman (bottom row) plots comparing unfiltered and filtered tract-specific g-ratio data based on tract caliber and representation across participants. The scan-rescan repeatability is high for large caliber tracts (middle column) and even higher in tracts present across at least 50% of participants (right column).

Figure 3 compares the scan-rescan repeatability of tractometry and tract-specific g-ratio results. Tractometry demonstrates a higher ICC than tract-specific g-ratio (ICC=0.932 and 0.735, respectively) and displays narrower limits of agreement in the Bland-Altman plot. This outcome is anticipated due to the blurring caused by partial volume effects, which increases repeatability across scans.

**Figure 3:**
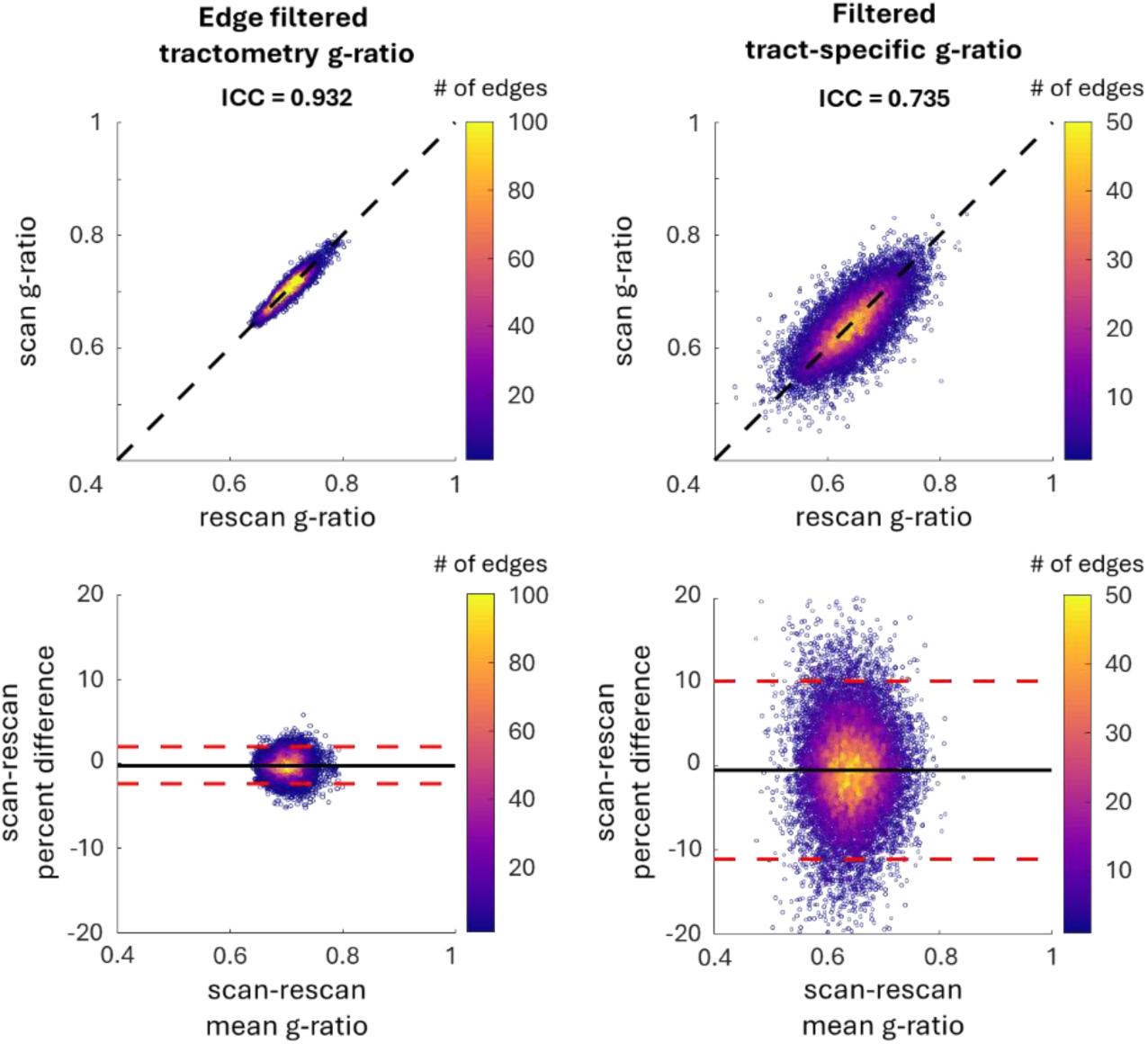
Comparison of filtered tractometry and filtered tract-specific g-ratio data. Correlation plots are in the top row and Bland-Altman plots in the bottom row. Tractometry exhibits an ICC of 0.932, with upper and lower limits of agreement in the Bland-Altman plot at 1.94% and -2.38%, respectively. Meanwhile, filtered tract-specific data demonstrates an ICC of 0.735, with limits of agreement at 10.1% and -11.2%, respectively.

### Tract-specific approach yields lower g-ratios and enhances contrast between tracts and networks compared to tractometry

While the overall topology of the g-ratio-annotated connectome is similar between the two techniques, there are also several differences. The most notable difference is that tract-specific g-ratio estimates are lower, corresponding to thicker myelin sheaths, compared to tractometry (Figure 4). When calibrating the MVF map using the ICVF maps from NODDI (tractometry) and COMMIT (tract-specific), the calibration factors α_*calib*_ differed slightly: 19.63 and 20.48, respectively. This difference in splenium ICVF and thus α_*calib*_ is due to the whole brain optimization used in COMMIT, which assumes the ICVF is constant along a streamline, in contrast to the voxel-based fitting perform by NODDI. Furthermore, the g-ratio histogram is broader for tract-specific estimates compared to tractometry, indicating a greater contrast due to removing partial volume effects. Further analysis examined the voxel-wise percentage difference between the g-ratio map derived from NODDI and the tract-specific volumetric g-ratio map computed using the AVF and MVF maps from COMMIT (Figure S3). While the percentage difference in voxels along major white matter pathways is low (0-5 %), it increases in areas with partial voluming with the cortex and subcortical regions.

**Figure 4:**
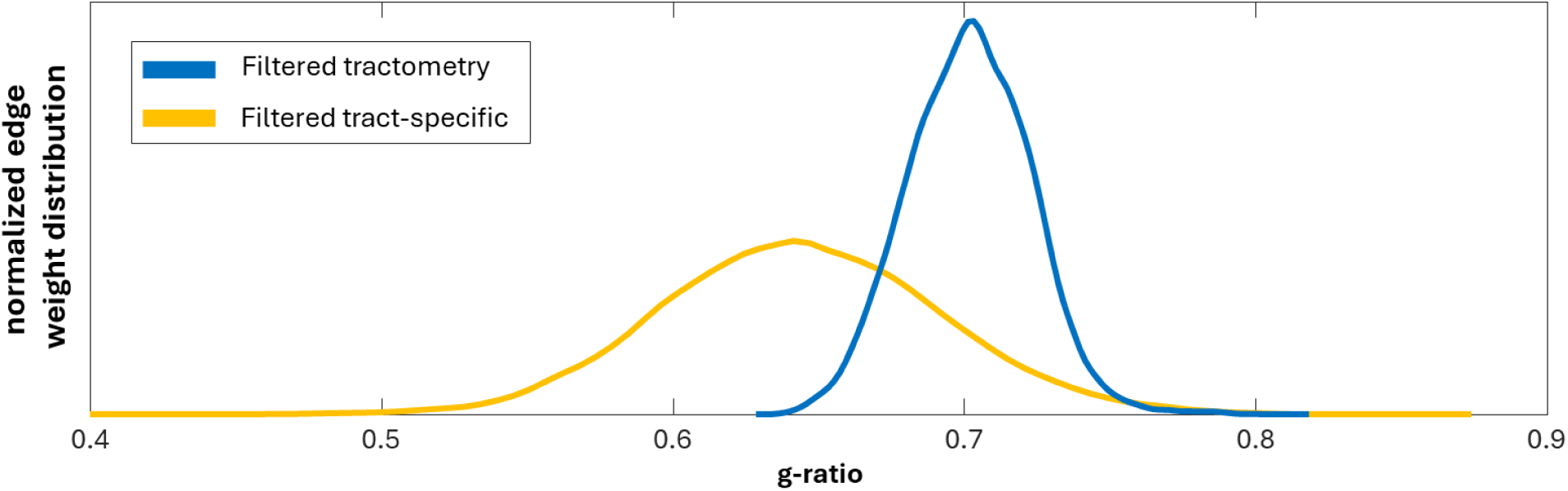
Normalized edge distribution of the g-ratio across all subjects. The edge-filtered tractometry is shown in blue, while the percentile and consensus-filtered tract-specific g-ratio is shown in yellow.

The data was z-scored to compare the topology of the two g-ratio annotated connectomes. The connections between the visual and salience/ventral attention nodes have the lowest g-ratio for tractrometry, while the connections between the somatomotor network and limbic are the highest (Figure 5). These trends differ from the tract-specific results, where the visual to limbic connections have the lowest network g-ratio, and the highest is observed in the subcortical-somatomotor network connections. Note that the absence of edges connecting the visual and somatomotor nodes does not imply a lack of connections between these two networks; rather, they were filtered out due to low tract caliber or consensus filtering.

**Figure 5:**
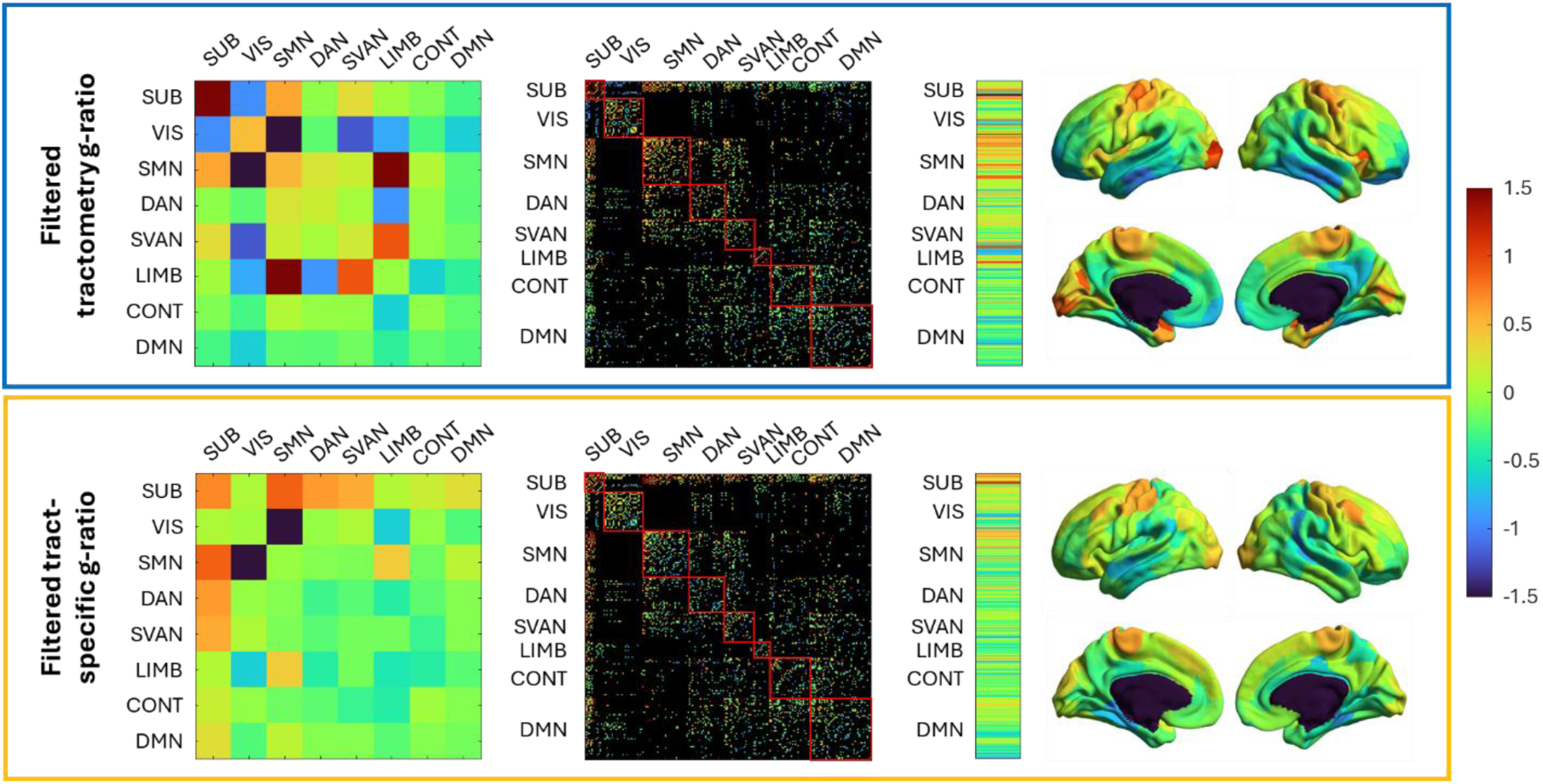
Comparison of z-scored g-ratio within resting-state network, connectivity matrices, and spatial patterns between tractometry and tract-specific. The top row (blue) displays the edge-matched tractometry g-ratio results, whereas the bottom row (yellow) displays the percentile and consensus filtered tract-specific results. The first column displays the mean g-ratio between resting-state functional networks: subcortical (SUB), visual (VIS), somatomotor (SMN), dorsal attention (DAN), salience/ventral attention (SVAN), limbic (LIMB), control (CONT), default mode network (DMN). The second column represents the g-ratio annotated connectivity matrices. The third column displays the node’s mean tract g-ratio across subject, with the results projected onto a cortical map.

The average g-ratio of white matter tracts connected to each cortical node was computed and projected onto the cortical surface to visualize the distribution of g-ratios across different cortical regions (Figure 5). The surface projections reveal different spatial patterns, with higher g-ratios (indicative of thinner myelin sheaths) in tracts connected to the visual cortex and lower g-ratios in those connected to the default mode network and frontal pole in tractometry compared to tract-specific. However, both techniques also show similar patterns: high g-ratio values in the motor regions and moderate g-ratio values in the parietal regions.

As in previous work (Leppert et al., 2023), we examine the crossing of the pontine fibers with the corticospinal tract (CST) in Figure 6. The blue box highlights voxels where these fibers exhibit varying degrees of partial volume. We specifically focus on voxels with a dominant single fiber orientation to better validate our g-ratio measurements and assess the impact of partial voluming on our results. This approach enables us to evaluate the accuracy of our g-ratio measurements in the absence of histological ground truth. The mean g-ratio of the pons, derived from the volumetric g-ratio map where voxel fractional anisotropy (FA)—which measures the degree of water diffusion in its primary direction—exceeds 0.5, is 0.679. In comparison, the g-ratio of the pons measured through tractometry is 0.718, suggesting partial voluming from tracts emerging from the brainstem. Conversely, the g-ratio of the pontine tract itself is 0.657, which aligns more closely with the volumetric map values and indicates less partial voluming.

**Figure 6:**
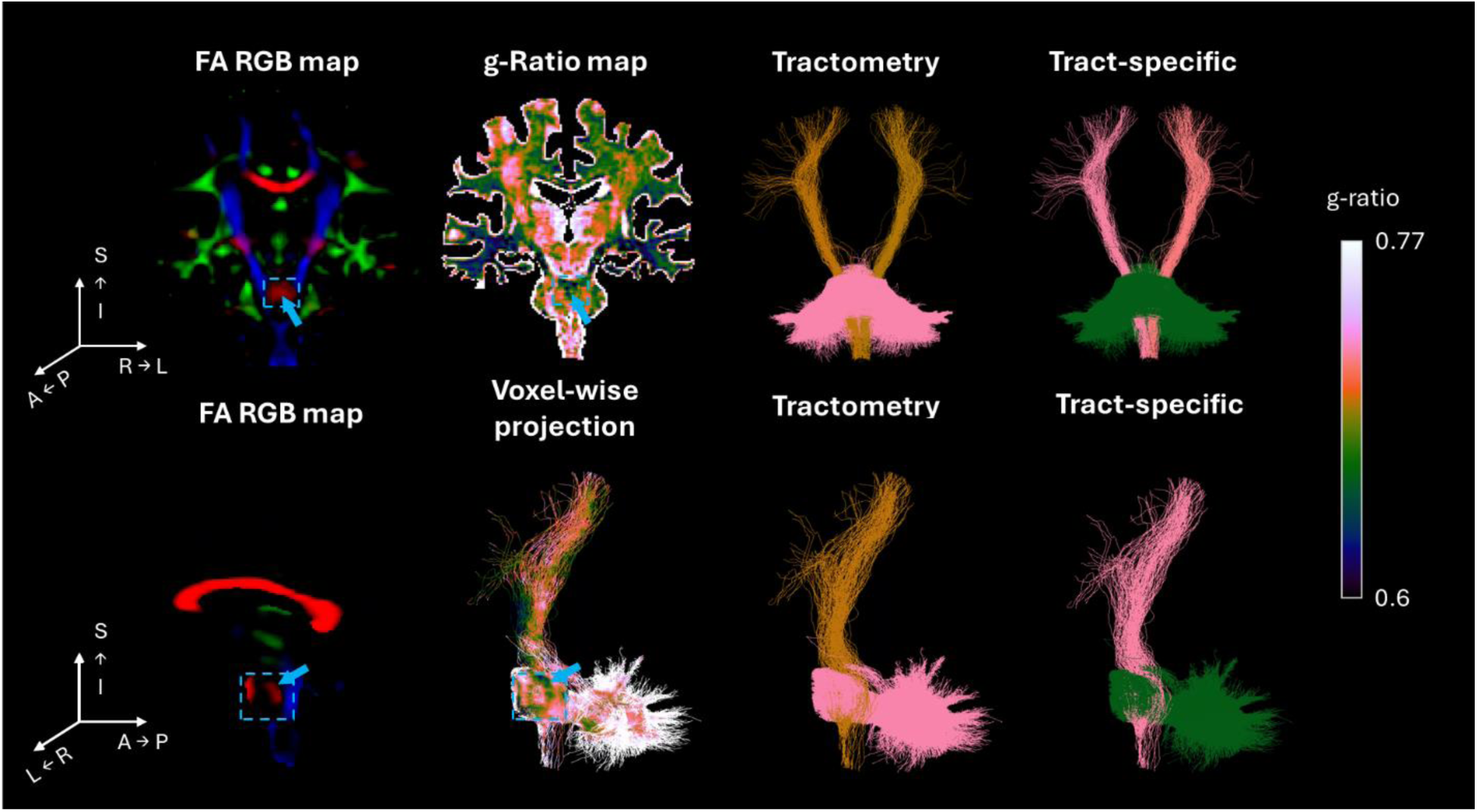
Comparison of tractometry vs tract-specific g-ratio at the intersection of the pons and cortico-spinal tract in a representative subject. Top row – coronal view: The left column displays the RGB FA map, with the arrow indicating the voxels traversed by the pons. The next column illustrates the aggregate g-ratio map, with the same arrow indicating the voxels traversed by the pons. The third column shows the pons and the brain-stem emerging tracts obtained from tractometry, and the fourth column shows the tract-specific approach. Bottom row – sagittal view: The left column represents the voxel wise g-ratio value projected onto the streamlines. The second column shows the pons and the brain-stem emerging tracts obtained from tractometry, and the third column shows the tract-specific approach. Data from additional subjects are included in Supplementary Material (Figure S4).

For the CST, the tract-specific g-ratio is 0.714, while the tractometry value is lower at 0.693. However, we cannot obtain a reliable g-ratio for the CST from the volumetric map due to partial voluming with other white matter tracts in the voxels it crosses. Interestingly, the contrast between the pons and CST is flipped between tractometry and tract-specific analysis, highlighting the differences between the two methods.

Figure 7 presents additional whole brain views of the participant tractogram shown in Figure 6. After z-scoring (top two rows), we observe that the tract with the highest g-ratio differs between the two methods: the CST (red arrows) has the highest g-ratio in the tract-specific analysis, while the pontine fibers (yellow arrows) have the highest g-ratio in tractometry. Despite this difference, both methods consistently show a high g-ratio value for the CST compared to connections to other cortical areas. Additionally, the g-ratios of tracts connecting to the frontal (blue arrows) and parietal (purple arrows) cortices, as well as the corpus callosum (green arrows), are significantly lower in tractometry compared to the tract-specific approach. The bottom two rows of Figure 7 display the same g-ratio-annotated tractogram without z-scoring, highlighting the broader distribution of g-ratio values for the tract-specific method. Tractograms of additional participants can be found in Figures S5-6 of the Supplementary Material.

**Figure 7:**
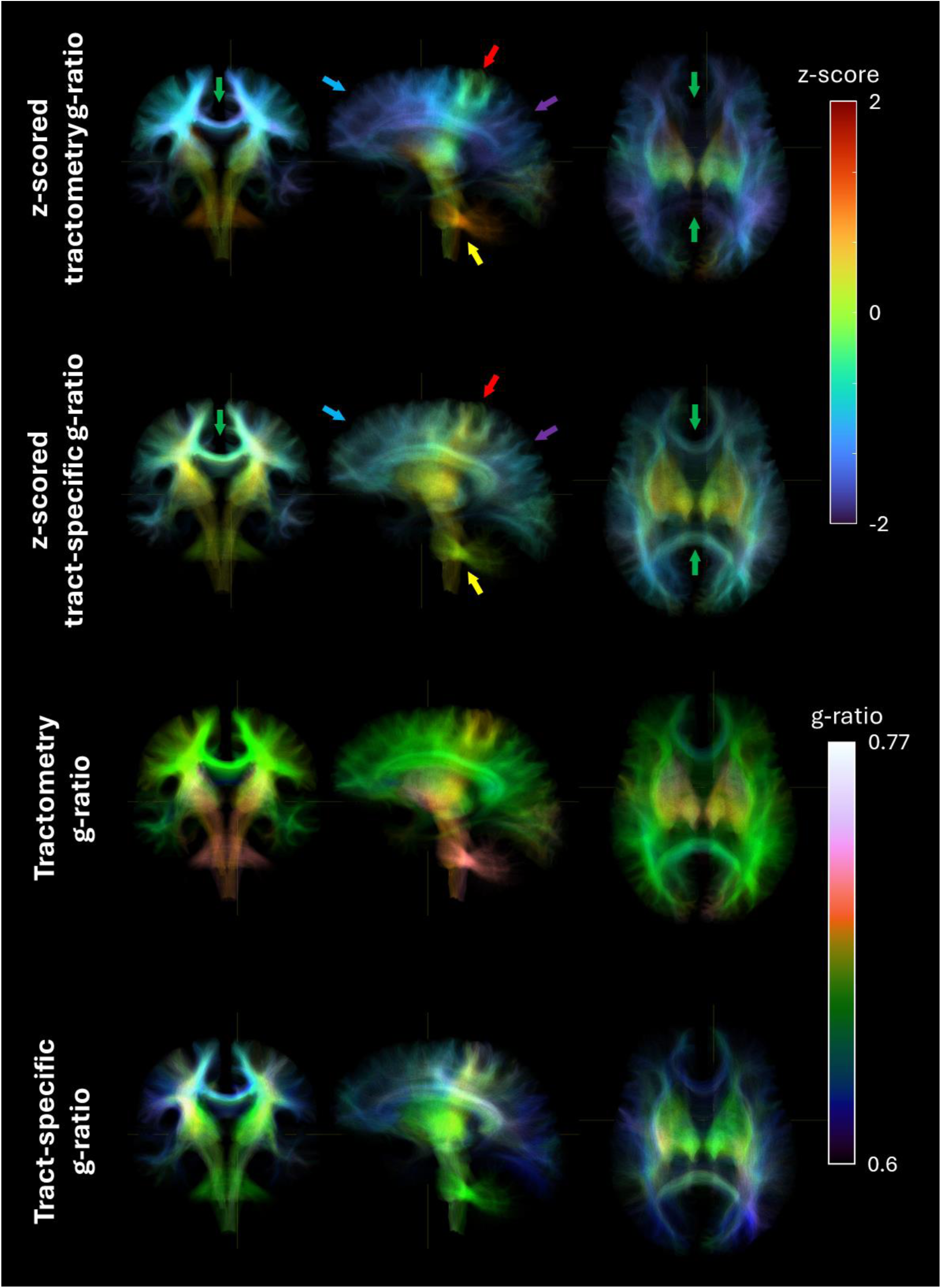
Streamlines spanning the entire brain of a representative subject. The top two rows are color-coded based on z-scored g-ratios, while the bottom two rows use g-ratio values. The arrows highlight key regions of interest: red arrows point to the CST, yellow arrow points to the pontine fibers, blue arrows indicate the frontal region, purple arrows highlight the parietal region, and green arrows mark the corpus callosum.

### Higher coefficient of variation across subjects for tract-specific in comparison to tractometry g-ratio results

The coefficients of variation (CoVa) of the tract g-ratio estimates across subjects for both techniques are shown in Figure 8. For all subsequent analyses, the edge values were averaged across scan and rescan for each subject. For both techniques, the intra-hemisphere variation is similar to the inter-hemisphere variation. At the resolution of edges, the CoVa for tractometry was lower than for tract-specific g-ratio measurements (Figure 8A), suggesting that tract-specific g-ratio is more sensitive to inter-individual differences. Regarding network connections, LIMB-LIMB and SUB-SUB edges exhibit the highest CoVa for tractometry, while VIS-SVAN has the lowest CoVa. Conversely, for tract-specific g-ratio, SMN-CONT displays the highest coefficient of variation, while the edges connecting the SMN-LIMB have the lowest CoVa (Figure 8B). The CoVa at the resolution of nodes (i.e. all edges connected to a cortical node across subjects) is higher for the tract-specific results. Tractometry shows a higher CoVa in the occipital lobe compared to the rest of the brain, whereas the tract-specific analysis reveals higher CoVa in the frontal nodes.

**Figure 8:**
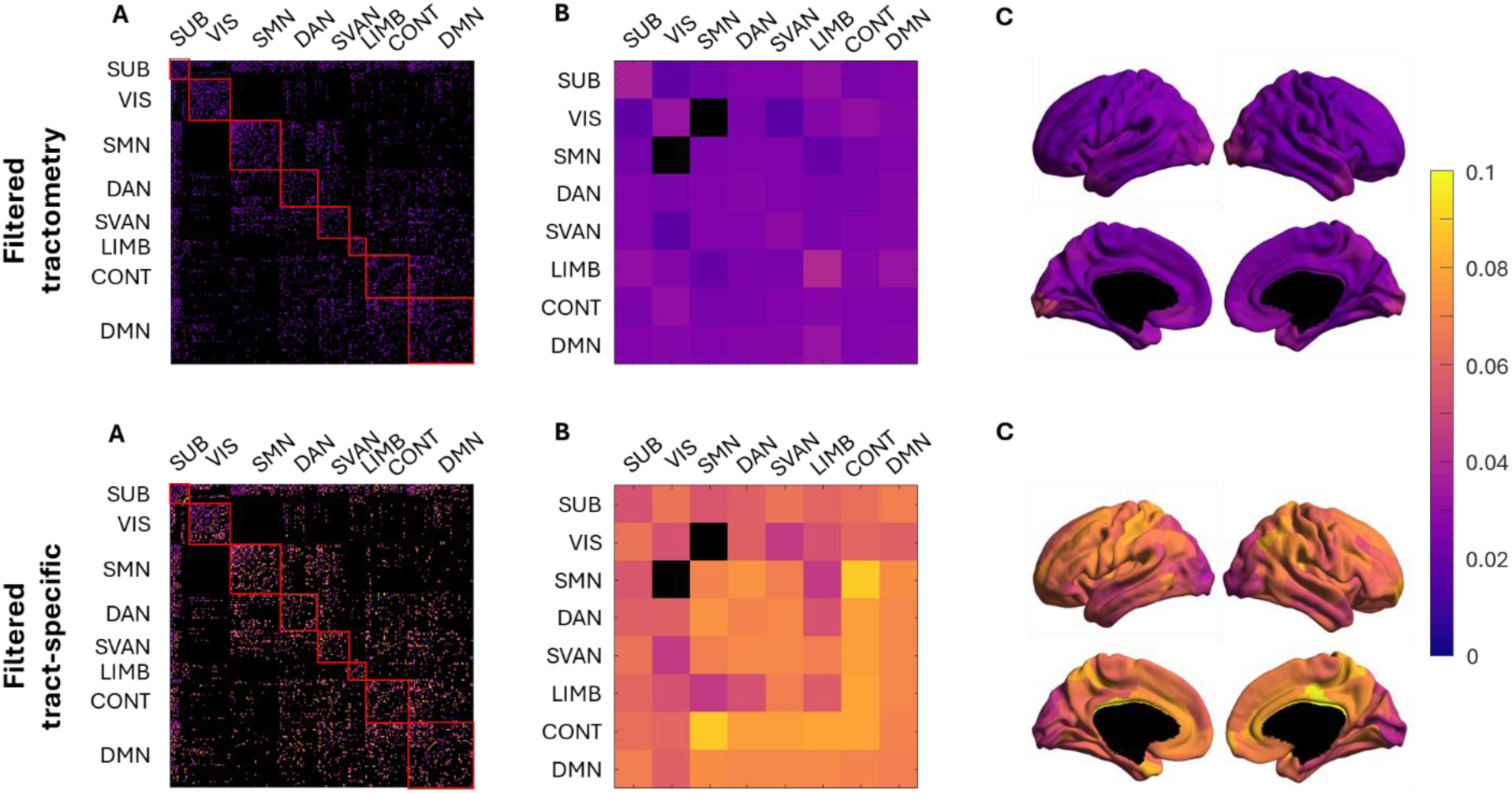
Coefficient of variation of tractometry and tract-specific g-ratio estimates across individuals (A) The CoVa of tractometry and tract-specific at the edge level is illustrated, with black edges indicating no tracts between nodes. (B) CoVa of edges connecting the resting-state functional networks, with black edges indicating no edges connecting the two networks. (C) CoVa of all edges connecting to a cortical node, projected onto the cortical surface.

### Tract-specific and tractometry g-ratio estimates are correlated

Given that several prior MRI-based g-ratio mapping studies have employed tractometry, it is crucial to evaluate whether significant discrepancies exist. In Figure 10 Figure, the correlation between tractometry and tract-specific results is depicted.

Inter-hemisphere edges show a higher correlation (0.529) between tractometry and tract-specific g-ratio measurements compared to intra-hemisphere edges (0.365). While most edges demonstrate a positive correlation between the two methods, a few edges have correlations that are close to zero or even negative. Edges connecting the SUB-VIS canonical networks have the lowest correlation but remain positive, while edges connecting LIMB-LIMB and SVAN-LIMB have the highest correlation between the two techniques. The projection onto the cortical surface of the correlation of all edges connected to a node is displayed in Figure 9C. Nodes at the occipital pole, isthmus cingulate cortex, and inferior frontal cortex exhibit lower correlations between the techniques, while those in the temporal and middle and superior frontal lobes show higher correlations.

**Figure 9:**
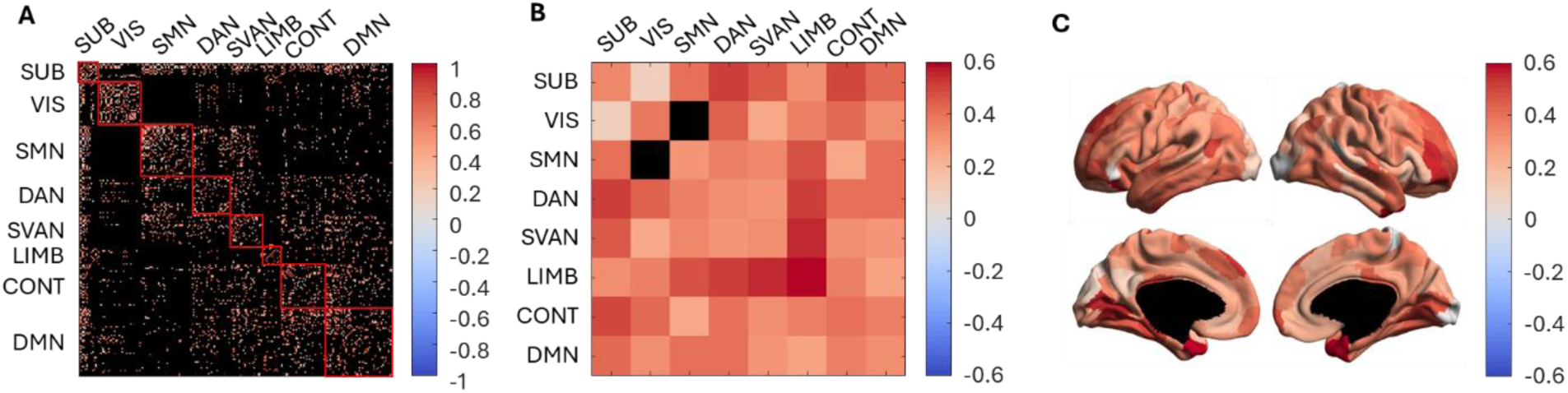
Correlation between the g-ratio estimated using tractometry and tract-specific pipelines. (A) The correlation between tractometry and tract-specific at the edge level is illustrated. (B) Correlation of edges connecting the functional networks, with black edges indicating no tracts between the two nodes. (C) Correlation of all edges connecting to a node across all subjects and projected onto the cortical surface.

### Tract-specific and tractometry g-ratio estimates exhibit opposite trends with tract length and caliber

The aggregate g-ratio is an intensive characteristic of a white matter tract, presumed to be uniform across the tract cross-sectional area and length. We explore the relationship between g-ratio and extensive tract attributes, length and caliber, across tracts in the brain. Since tract caliber is influenced by node size, it has been scaled by the inverse of the sum of the two node volumes. We used a mixed linear effects model to investigate the fixed effects for caliber, length, and length squared, as well as random effects for subject ID and for length and length squared grouped by subject ID:

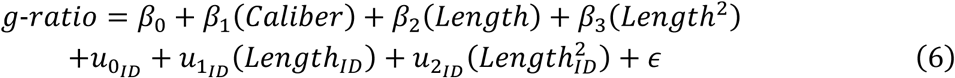

Figure 10 depicts the relationship of the g-ratio with tract caliber (above) and tract length (below), for both tractometry and tract-specific methods. For both tractometry and tract-specific g-ratio, the effects of caliber, length, and length squared were statistically significant (p< 0.001). However, the model fit was better for tractometry (adjusted R^2^=0.546) in comparison to the tract-specific (adjusted R²=0.186).

**Figure 10:**
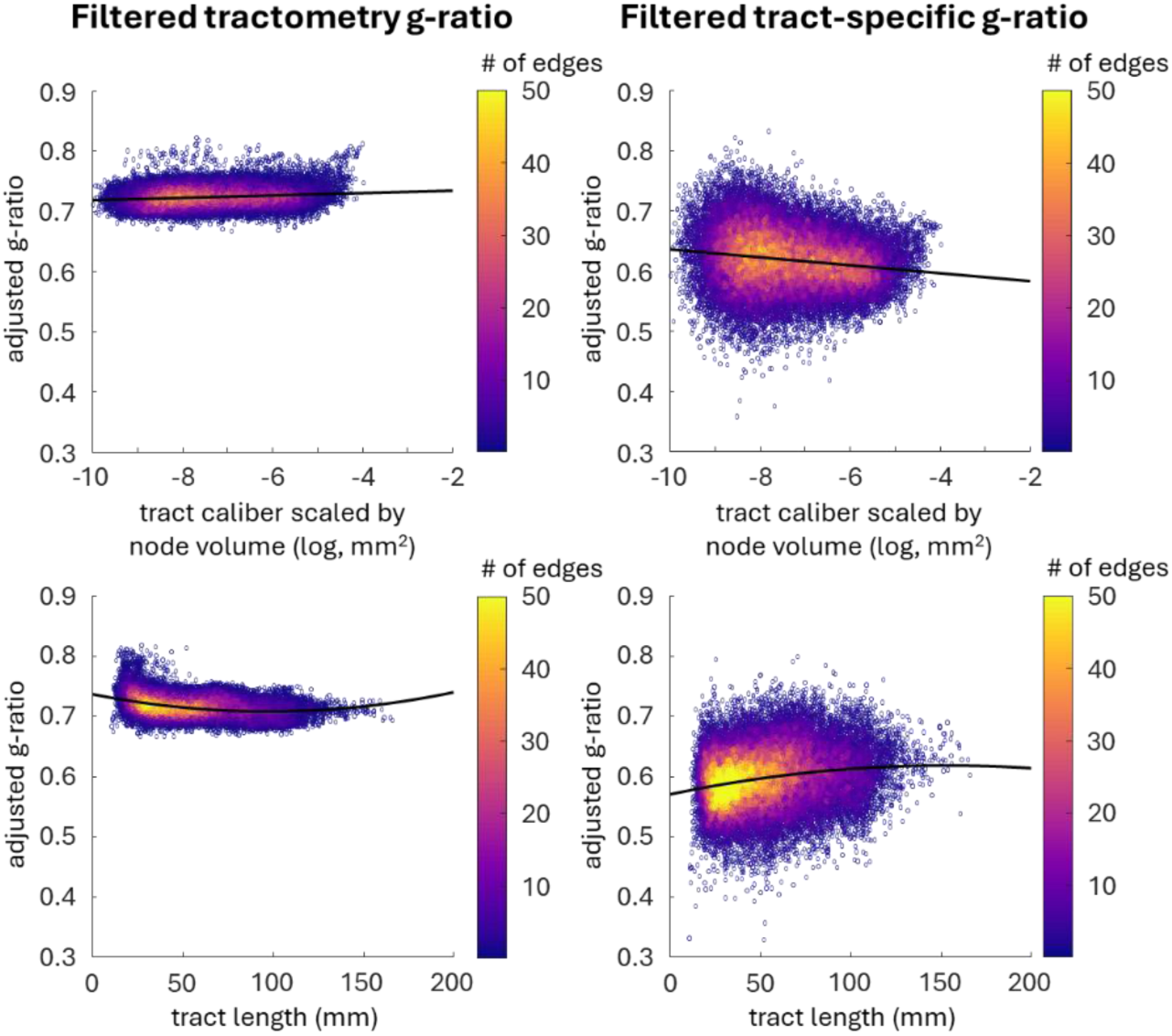
Relationship between g-ratio and tract caliber (top) and length (bottom). The adjusted g-ratio corresponds to the g-ratio with the random effect and the other fixed effects removed. Tractometry results are displayed on the left, and tract-specific results on the right. The black line in each plot represents the line of best fit. Tract-specific results are accompanied by the trend of a broader distribution of g-ratio.

In the tract-specific analysis, the model predicts a negative β_1_, suggesting that larger caliber tracts tend to have lower g-ratios, i.e. thicker myelin sheaths relative to axon diameter. Conversely, tractometry reveals the opposite trend, with a positive β_1_, indicating that g-ratios increase as tract caliber increases.

When examining the relationship between g-ratio and tract length, both methods again show opposing patterns. For tract-specific g-ratio, the model predicts a positive β_2_and a negative β_3_, producing an inverted U-shaped curve, where g-ratio increases as tract length increases. In contrast, tractometry predicts a negative β_2_and a positive β_3_, following a U-shaped curve, where g-ratios decrease until ∼95 mm in length, after which they begin to increase. These findings highlight fundamental differences in how each method captures the relationship between g-ratio and the structural characteristics of the tracts.

## 4 Discussion

Tractometry, the current state-of-the-art, has limited anatomical specificity (Schiavi et al., 2022). It samples each voxel traversed by a streamline, then takes the mean or median to estimate the streamline’s g-ratio. Most of the traversed voxels contain multiple fiber populations, each with specific microstructural properties, which leads to partial volume effects that are compounded along the length of the streamline. This introduces a bias and reduces the anatomical specificity of tract g-ratio estimates obtained from tractometry. To address this issue, this study expands on the COMMIT framework, previously used to derive tract-specific intra-axonal and myelin volumes, to estimate the tract-specific aggregate g-ratio. Since the g-ratio is not additive, the COMMIT framework cannot be applied directly to the g-ratio map. Instead, the g-ratio is computed at the bundle level using MV and AV estimates derived from MTsat and DWI data, respectively using COMMIT. This approach effectively mitigates the partial volume effects caused by crossing fibers.

### Repeatable tract-specific g-ratio mapping of large caliber tracts

The tract-specific g-ratio estimates are highly repeatable across scanning sessions in large tracts, as evidenced by an ICC of 0.715 at the Schaefer-200 resolution. However, when looking at all tracts, the ICC decreases considerably to 0.175, indicating that the g-ratio estimates of small caliber tracts are less repeatable. The axon caliber and myelin caliber estimates for these smaller tracts are especially susceptible to error from noise, which propagates to the g-ratio calculation and significantly increases its variance. Applying a 50% edge consensus filter across participants increases the confidence of the remaining tracts, and further increases the ICC to 0.727. The tract-specific g-ratio pipeline can be applied to higher parcellation resolutions (i.e., resulting in smaller caliber tracts), albeit with slightly reduced ICC values (e.g., 0.611 for the filtered Schaefer-400 results). The first iteration of COMMIT filters the streamlines that do not fit the model to the DWI data to remove false positive streamlines. However, the technique is unable to eliminate all false positives and cannot correct for false negatives, its effectiveness relies heavily on the quality of the input tractogram (Schiavi et al., 2020). Therefore, using a more accurate tractogram can improve the repeatability of the technique, especially for smaller tracts.

### Improved anatomical specificity results in enhanced g-ratio contrast between tracts and individuals

Tract-specific and tractometry aggregate g-ratio estimates align closely with previous neuroimaging literature examining whole-brain voxel-wise aggregate g-ratio estimates (Mohammadi et al., 2015), typically ranging from 0.5 to 0.8. Our tract-specific g-ratio estimates tend to be lower than those observed in tractometry, accompanied by a much broader distribution of g-ratio values. Additionally, the coefficient of variation across subjects for tract-specific g-ratio is much higher than that of tractometry. This difference is likely attributed to the partial volume effect observed along the length of the tracts in tractometry, which also contributes to an increased scan-rescan repeatability compared to the tract-specific technique. The coefficients of variation in g-ratio tractometry are mostly below 0.3, in line with previous g-ratio research (Cercignani et al., 2017; Mohammadi et al., 2015). Whereas for tract-specific g-ratio, the variability between participants tends to be much more pronounced, a trend seen in previous tract-specific microstructure mapping studies (Leppert et al., 2023; Nelson et al., 2023). The increased dynamic range observed in tract-specific results has also been observed in prior studies that address partial voluming (De Santis et al., 2016; Leppert et al., 2021; Leppert et al., 2023; Schiavi et al., 2022).

Given that prior MRI studies on g-ratio have employed tractometry (Cercignani et al., 2017; Mancini et al., 2018; Slater et al., 2019), we conducted a correlation analysis between tractometry and tract-specific g-ratio results to determine the level of agreement or disagreement between the two techniques. When examining inter- and intra-hemisphere results, the correlation between tractometry and tract-specific g-ratio values is positive (0.37 for intra-hemisphere and 0.53 for inter-hemisphere), indicating a general agreement between the two approaches. While most edges from the Schaefer-200 parcellation exhibit a positive correlation, there are instances where the correlation approaches zero or is negative. This suggests that certain tracts may traverse one or several other tracts such that their individual tract characteristics are significantly biased due to partial volume effects.

### Tract-specific and tractometry reveal opposite relationships between the g-ratio and tract attributes

When examining the g-ratio correlation across subjects, the trend is generally positive: both tractometry and tract-specific analysis agree on whether a particular subject’s g-ratio is higher or lower compared to others. However, when we explored the relationship of tract g-ratio with tract length and caliber, we found contrasting patterns between the two techniques. Tractometry shows a positive effect of tract caliber on g-ratio estimates, a negative effect of length, and a positive effect of length squared. In contrast, tract-specific analysis reveals the opposite polarity: a negative effect of tract caliber on the g-ratio, a positive effect of tract length, and a negative effect of length squared. This suggests that the agreement between tractometry and tract-specific analysis holds only at the broader, cross-subject level. But when we delve deeper into specific tract attributes, this agreement is weaker, and the differences between the two techniques become more pronounced.

Previous rabbit histological studies of the splenium (Waxman & Swadlow, 1976) reported no significant correlation between g-ratio and tract caliber. In our study, we similarly found no relationship in the splenium (data not shown). In contrast, the whole-brain analysis shows a significant effect of both caliber and length on tract g-ratio. However, the model’s low adjusted R² indicates that while the relationships are statistically significant, length and caliber explain only a limited portion of the variance in the g-ratio. This suggests that other factors, such as age, gender, genetics, and environmental influences, may modulate myelination and g-ratio.

There is a noticeable shift between the g-ratio values obtained from tractometry and tract-specific methods, with tract-specific estimates being lower, indicating thicker myelin sheaths. This discrepancy likely arises from differences in how COMMIT and NODDI handle partial voluming with gray matter. Additionally, the resolution mismatch between DWI (2.6mm) and MTsat (1mm) images—MTsat typically offering higher resolution than DWI—may introduce interpolation artifacts in voxels near the cortex and subcortical grey matter. These factors influence the voxel’s ICVF and, consequently, the g-ratio calculation.

When examining the relationship between g-ratio and tract length, distinct trends emerge between the two techniques. Tractometry shows a relatively constant g-ratio as tract length increases, while tract-specific g-ratio values rise with increasing tract length. The volumetric percentage difference map between the two techniques reveals 0-5% differences in g-ratio of major white matter pathways, indicating a good agreement for longer tracts. However, larger differences are found near the cortex and in subcortical regions, suggesting that these areas contribute to the overall shift in g-ratio distribution. Taken together, these observations imply that part of the difference between the two techniques stems from tracts traversing subcortical regions and shorter white matter fibers, such as short-range association or U-fibers.

### Limitations and future work

The tract-specific aggregate g-ratio mapping technique was developed to improve anatomical specificity in comparison to conventional tractometry. There is unfortunately no ground truth from large scale histological studies to validate our findings. We therefore compared the tract-specific results to voxels that contain a single fiber as a form of validation.

Our tract-specific g-ratio mapping method uses the COMMIT framework to first disentangle the axonal and myelin volumes of crossing tracts in the brain. We are thus limited by the assumption that the microstructural properties of these tracts are constant along their length. This assumption may not always hold true, particularly in pathology such as focal white matter lesions. Ongoing work is focused on adapting the framework to account for these lesions and integrate them into the analysis (Bosticardo et al., 2023).

The current edge or tract filtering method employed is quite aggressive, effectively removing smaller tracts to reduce noise. This eliminates all edges between certain networks, specifically between the VIS and SMN networks. This highlights a significant challenge in balancing the need to filter out false positives with the risk of losing genuine connections.

It is well known that MRI-based tractography has inherent limitations, particularly in generating false positive streamlines (Maier-Hein et al., 2017). Determining the appropriate matrix density remains a point of contention due to the lack of consensus and absence of a definitive ground truth (Schiavi et al., 2020). This underscores the complexity of filtering streamlines and edges to achieve both accuracy and completeness in structural network connectivity.

Tract-specific g-ratio mapping can be carried out using widely available MRI techniques. In this study, MTsat was employed to evaluate tract myelin volume, but other myelin-sensitive methods such as myelin water imaging (Schiavi et al., 2022) or inhomogeneous magnetization transfer (Berg et al., 2022) are also effective alternatives. There are however two main limitations in modeling the ICVF. First, both the NODDI and the COMMIT model fix diffusivities. While these values can vary slightly between tracts (Vos et al., 2012), they also change with age (Hasan et al., 2014) or in the presence of pathology (Chung et al., 2017). Implementing the standard model (Novikov et al., 2019) into COMMIT for diffusivity estimation could help address this issue. Second, NODDI and the StickZeppelinBall model used in COMMIT output T2-weighted signal fractions for the stick compartment, but we have treated them as volume fractions (as is commonly the practice). These models do not account for compartment-specific T2 relaxation times (Papazoglou et al., 2024) which can bias g-ratio estimates (Gong et al., 2020). This limitation becomes more pronounced in pathological conditions, where T2 times can significantly differ from those in healthy white matter (UDAKA et al., 2002). To accurately measure compartment volume fractions, a diffusion-relaxometry acquisition with multiple echo times is needed (Barakovic et al., 2021).

Tract-specific g-ratio mapping could contribute to a better understanding of brain network microstructure during neurodevelopment, aging, and in disease. Using g-ratio annotated connectivity matrices, the impact of myelin g-ratio on the relationship between network structure and function can be studied, providing a more comprehensive understanding of brain network organization and dynamics. This technique holds potential to study the impact of adaptive myelination on brain network function (Knowles et al., 2022), and can provide insights into the characteristics of white matter tracts and networks in various disorders such as autism or Alzheimer’s disease.

## 5 Conclusion

This study presents a novel method for calculating the aggregate g-ratio of individual white matter tracts using the COMMIT framework, aimed at improving anatomical specificity. By filtering out false positive streamlines and small-caliber edges, the approach ensures repeatable g-ratio estimates and enhances contrast between tracts and individuals compared to tractometry. Validation of this method in the pons, using a volumetric map, demonstrated that the tract-specific results closely matched the volumetric g-ratio measurements, reinforcing the accuracy of the approach. This technique advances tract-specific analysis by reducing biases from the complex network of crossing white matter fibers and demonstrates that integrating microstructural data further improves the potential of connectomics research.

## 6 Data and Code Availability

The full pipeline is available on Github: https://github.com/TardifLab/mwcpipe

## 7 Author Contributions

W.L.: writing, idea implementation and data analysis. M.C.N.: analysis and visualization code, image acquisition and manuscript editing. I.L.R.: image acquisition and manuscript editing. J.S.W.C.: discussion and manuscript edit. S.S.: support for COMMIT framework and manuscript editing. G.B.P.: discussion and manuscript edit. C.D.R.: sequence implementation and manuscript editing. A.D.: support for COMMIT framework and manuscript editing. C.L.T.: idea, financial support, manuscript editing and supervision.

## 9 Acknowledgements

This work was supported by the following funding sources: Canadian Neurodevelopmental Research Training Platform (CanNRT), Natural Sciences and Engineering Research Council of Canada (NSERC), Brain Canada, Fonds de Recherche du Québec— Santé (FRQS), and Healthy Brains Healthy Lives, Killam Trusts.

## 10 Supplementary Material

Supplementary information is available in an additional document.

