## Supplemental Material for "Mapping the aggregate g-ratio of white matter tracts using multi-modal MRI"

### Supplementary Material

Wen Da Lu[1,2], Mark C. Nelson[2,3], Ilana R. Leppert[2], Jennifer S.W. Campbell [2], Simona Schiavi[4], G. Bruce Pike [5], Christopher D. Rowley[6], Alessandro Daducci [4], Christine L. Tardif[1,2,3]

1. Department of Biomedical Engineering, McGill University, Montreal, QC, Canada
2. McConnell Brain Imaging Centre, Montreal Neurological Institute and Hospital, Montreal, QC, Canada
3. Department of Neurology and Neurosurgery, McGill University, Montreal, QC, Canada
4. Department of Computer Science, University of Verona, Verona, Italy
5. Hotchkiss Brain Institute, Department of Radiology, and Department of Clinical Neuroscience, University of Calgary, Calgary, AB, Canada
6. Department of Physics and Astronomy, McMaster University, Hamilton, ON, Canada

Corresponding authors:

Wen Da Lu,

Christine L. Tardif,

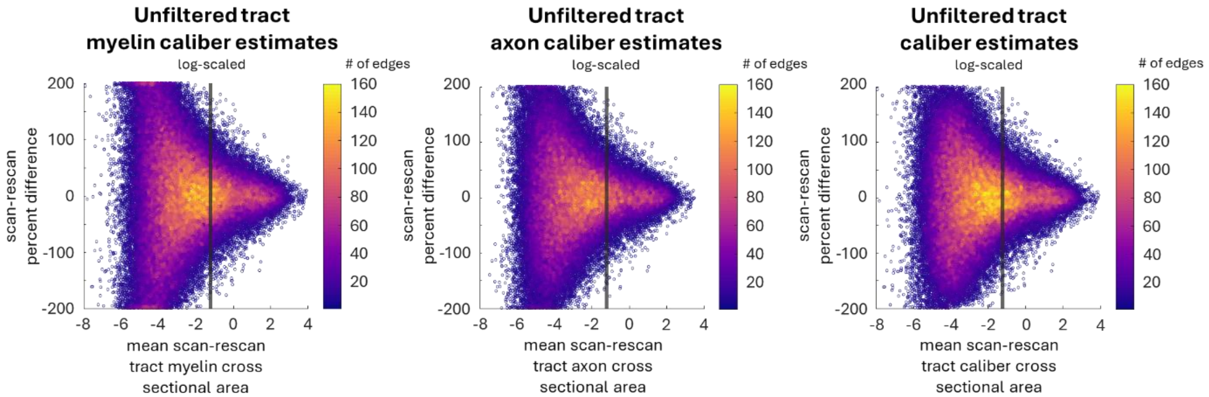


**Figure S1: Percent scan-rescan difference as a function of the tract's total myelin cross-sectional area, total axonal cross-sectional area, and tract caliber, i.e. the sum of the myelin and axonal areas (log-scaled).** The black vertical line indicates the 80% cutoff threshold used in the subsequent analyses. Larger tracts are more repeatable than for smaller caliber tracts.

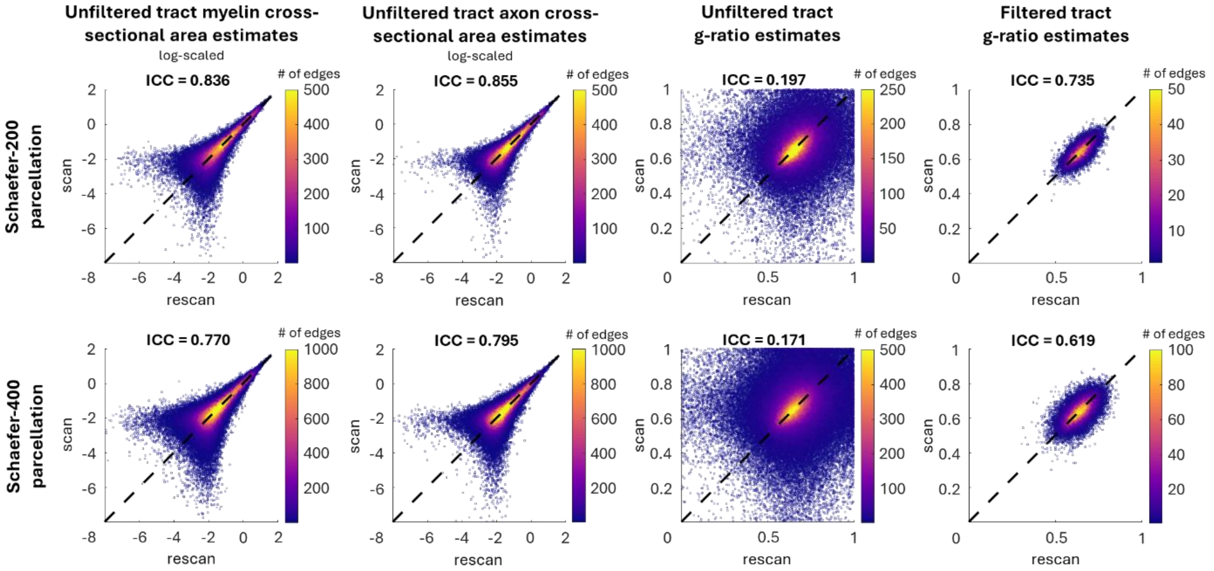


**Figure S2: Impact of cortical parcellation resolution on scan-rescan repeatability of tract-specific g-ratio.** The figure illustrates a tract's myelin caliber, axon caliber, unfiltered, and filtered g-ratio plotted against its rescan at two resolutions: Schaefer-200 (top row) and Schaefer-400 (bottom row). The correlation plot of myelin caliber and axon caliber has been log-scaled. The ICC for myelin caliber estimates at Schaefer-200 and-400 parcellations is 0.836 and 0.770, respectively. For the axon caliber estimates, the ICC at Schaefer-200 and-400 parcellations is 0.855 and 0.795, respectively. The ICC for unfiltered g-ratio estimates at Schaefer-200 and-400 parcellations is 0.197 and 0.171, respectively. Finally, the ICC for percentile and consensus filtered g-ratio estimates at Schaefer-200 and-400 parcellations is 0.735 and 0.619, respectively.

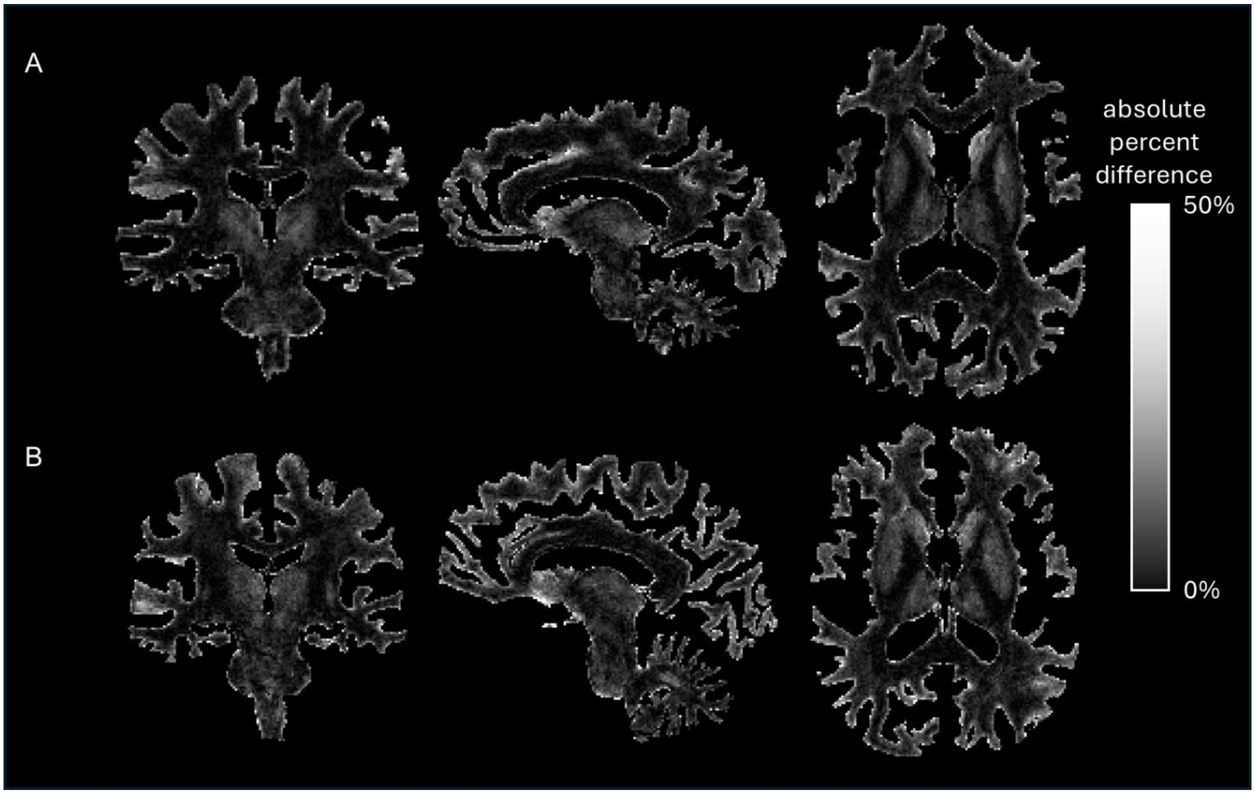


**Figure S3: The absolute percent difference between the g-ratio map computed from COMMIT's AVF and MVF maps and the volumetric g-ratio calculated from NODDI for two subjects**. While the percentage difference in voxels along major white matter pathways is low, it increases near the cortex and in the subcortical regions. This suggests that shorter white matter tracts and tracts traversing subcortical regions are the primary drivers of the g-ratio differences between tract-specific and tractometry approaches.

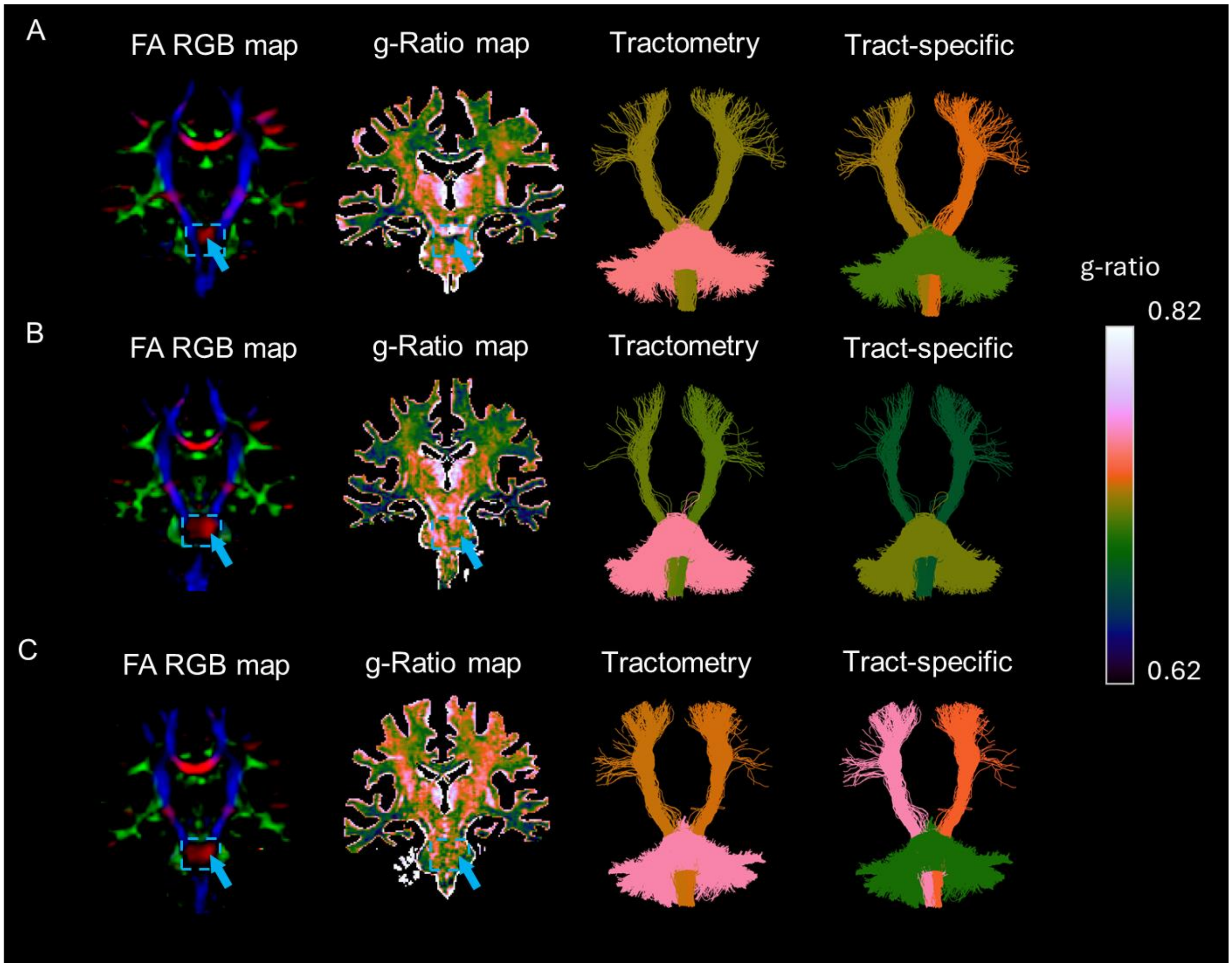


**Figure S4: Comparison of tractometry vs tract-specific aggregate g-ratio mapping of the pontine fibers and the corticospinal tract in three additional participants.** The left column displays the RGB FA map with an arrow indicating the voxels traversed by the pons (left-right direction in red). The second column illustrates the volumetric aggregate g-ratio map, with the same arrow indicating the voxels traversed by the pons. The third and fourth columns present the aggregate g-ratio of the pons and the tracts emerging from the brain stem obtained using tractometry and the tract-specific approach, respectively. The mean g-ratio of the pons derived from the volumetric g-ratio map is 0.729, 0.729, and 0.716 for subject A, B, and C, respectively in comparison to 0.752, 0.756 and 0.758 for tractrometry and 0.709, 0.719 and 0.700 for the tract-specific approach.

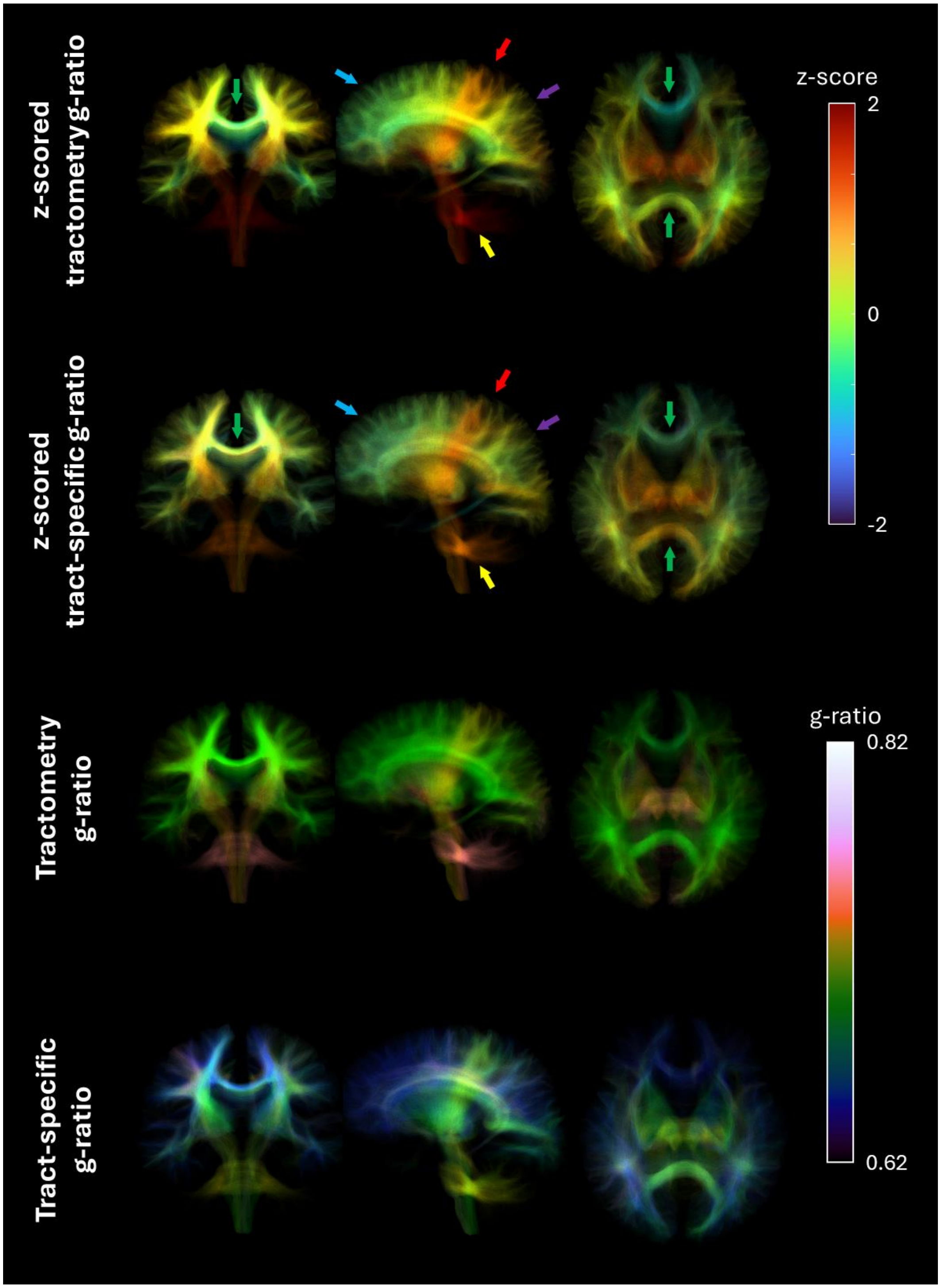


Figure S5: Streamlines spanning the entire brain of participant A from Figure S4. The top two rows are color-coded based on z-scored g-ratios, while the bottom two rows use g-ratio values. The arrows highlight regions of interest: red arrows point to the CST, where z-scored values are slightly higher in the tract-specific method; yellow arrows point to the pontine fibers, which show higher z-scored values in tractometry; blue arrows indicate the frontal region and purple arrows highlight the parietal region, where both methods yield similar z-scored values; and green arrows mark the corpus callosum, where z-scored values are lower in tractometry.

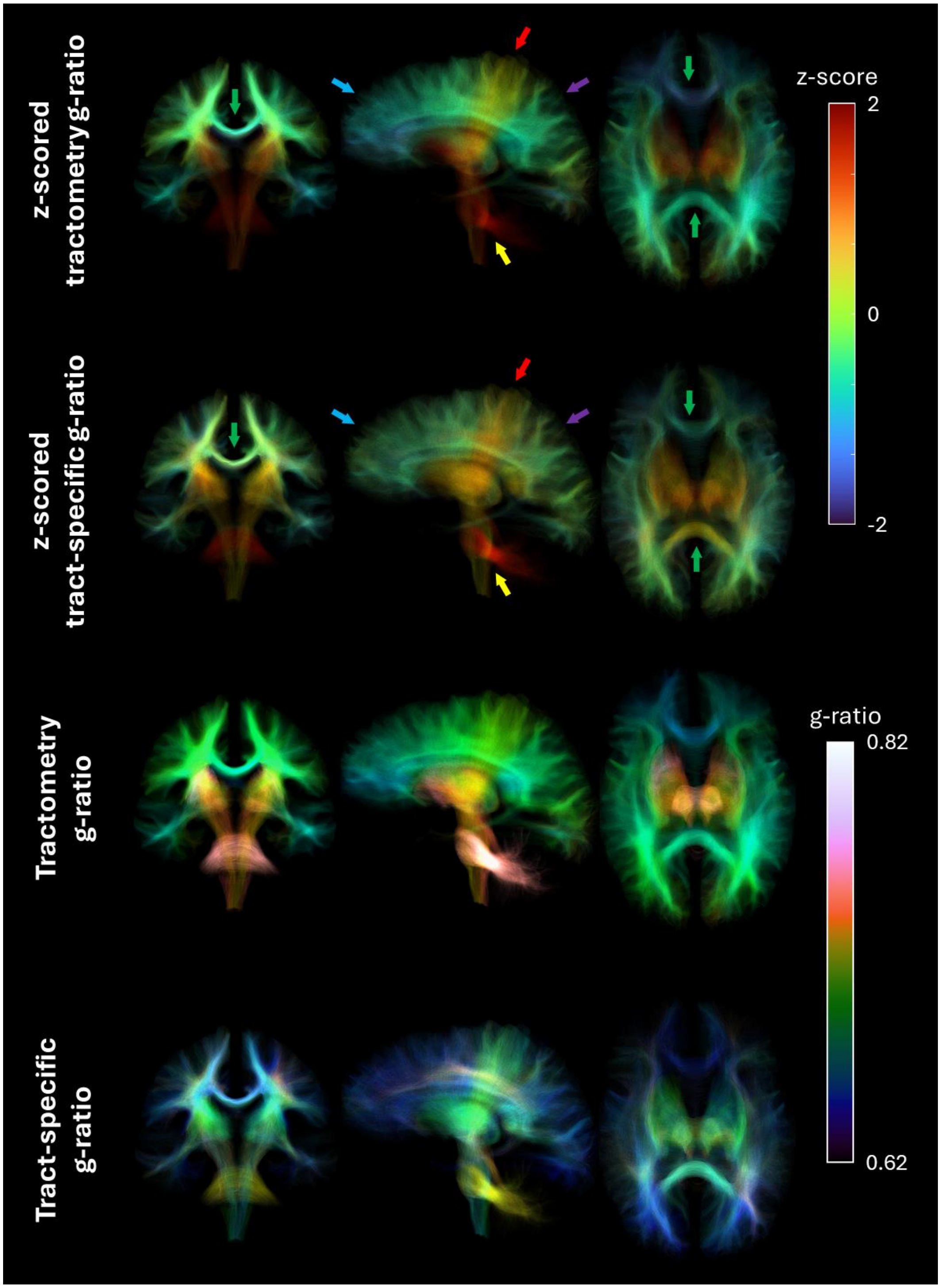

Figure S6: Streamlines spanning the entire brain of participant B from Figure S4.
